# *bifrost*: an R package for scalable inference of phylogenetic shifts in multivariate evolutionary dynamics

**DOI:** 10.64898/2026.04.12.718036

**Authors:** Jacob S. Berv, Nathan Fox, Matt J. Thorstensen, Henry Lloyd-Laney, Emily M. Troyer, Rafael A. Rivero-Vega, Stephen A. Smith, Matt Friedman, David F. Fouhey, Brian C. Weeks

## Abstract

1. High-dimensional comparative datasets, including geometric morphometric landmarks, functional traits, and other large trait datasets, are increasingly common in biology. When these datasets include a large number of traits relative to the number of taxa, they pose significant challenges for phylogenetic comparative analysis. In addition, evolutionary dynamics are often heterogeneous across phylogenies, challenging researchers to develop tools that can localize and account for such variation when investigating hypotheses of evolutionary change.
2. We present *bifrost*, an R package for detecting and characterizing shifts in multivariate trait evolution across phylogenetic trees. *bifrost* implements a stepwise greedy search over alternative macroevolutionary regime configurations on a phylogeny. Candidate shifts are proposed and assessed at internal nodes, accelerated with parallel model fitting where possible, and aggregated sequentially when they exceed a user-defined information-criterion acceptance threshold.
3. The underlying model is a scalar-rate multivariate Brownian motion process fit by generalized least squares using mvMORPH::mvgls [1]. Our framework also provides support estimates for individual shifts using information-criterion weights.
4. We illustrate the workflow using a fossil-tip-dated phylogeny and high-dimensional landmark data for early bony fish jaws (32,508 scalar coordinate values), and discuss tuning, outputs, and limitations. *bifrost* extends existing phylogenetic comparative frameworks for evolutionary analysis and provides a scalable pipeline for exploring the phylogenetic natural history of large multivariate datasets.

## Introduction

Although morphology has long been central to evolutionary biology and paleontology, generating and analyzing multivariate datasets remains a major challenge [2]. Traditional studies typically rely on low-dimensional, manually collected measurements (< 10^1^) —but phenotypes are inherently multivariate, shaped by developmental, functional, and evolutionary constraints that cannot be captured by limited sampling [3-6]. In many empirical systems, the traits that best capture biological form and function are high-dimensional vectors: geometric morphometric landmarks, shapes represented by semilandmarks, multivariate functional traits, or large collections of measurements. These data naturally encode both the magnitude of evolutionary change and the covariance structure among traits, which together describe how phenotypes explore morphospace through time [1, 4, 5].

Recent advances in imaging and automated feature extraction are beginning to enable high-throughput recovery of such morphological signals from museum specimens (∼10^3^–10^4^) [2, 7-9]. Yet, many commonly used evolutionary models are poorly suited to interrogating rich, high-dimensional datasets without first applying dimensionality-reduction techniques such as principal component analysis (PCA). Many comparative workflows proceed with a small set of components—a lossy procedure that can obscure evolutionary signals, distort trait covariances, and bias estimates of evolutionary processes [1, 6, 10-14]. These challenges mirror findings in computer vision and natural language processing, where models that operate directly on high-dimensional data consistently outperform those that rely on compressed or preprocessed features [15, 16]. Analytical approaches that avoid lossy preprocessing steps may thus be expected to outperform methods that rely on feature compression [17]. As a result, realizing the potential of high-dimensional comparative data hinges on developing statistical methods that can operate at the scale, complexity, and resolution at which such variation naturally occurs.

Another recurring challenge in the study of trait evolution is heterogeneity in patterns and processes of diversification. Rates of phenotypic evolution can vary across lineages, and entire clades may experience accelerated or decelerated evolution [18-20]. A central inferential task is therefore to identify where along a phylogeny evolutionary dynamics and/or patterns of evolvability might change [21-23], and to understand how those changes may affect the evolutionary process. Existing software packages employ a wide range of techniques, from stepwise model selection to Bayesian reversible-jump MCMC models (Table 1) (e.g., [18, 24-27]). However, most existing approaches are built around univariate assumptions and expectations, and inference can become intractable or unidentifiable when the number of traits assessed is large relative to the number of taxa [1, 10, 14].

**Table 1.**
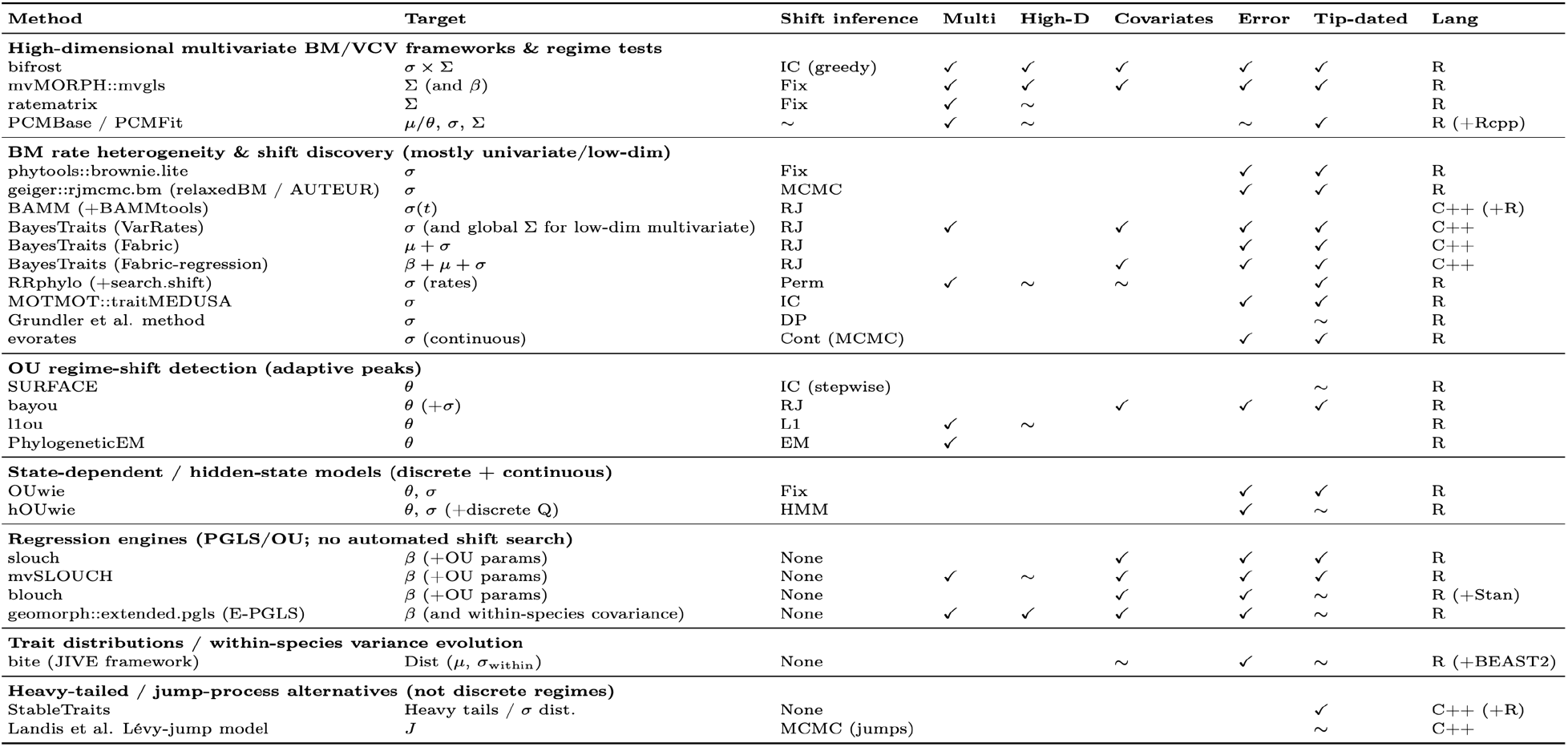
Software tools for phylogenetic analysis of heterogeneous continuous‐trait evolution. Methods are grouped by modeling paradigm (e.g., global tempo/mode models, fixed‐regime hypothesis tests, OU mean/optimum shift discovery, BM rate heterogeneity, episodic direction/evolvability models, mixed Gaussian/mixture frameworks, distributional trait evolution, and high‐dimensional GLS engines). Columns summarize (i) the primary evolutionary quantity that varies (μ = mean/optimum or directional dynamics; σ = rate/variance; Σ = multivariate evolutionary covariance; σ×Σ = proportional scaling of a baseline covariance; T = global tempo/mode; J = jumps/heavy tails; Mix = model mixtures), (ii) whether shifts/regimes are inferred versus user‐specified and the main inference strategy, and (iii) key practical capabilities relevant to high‐dimensional comparative workflows (multivariate and high‐dimensional support, regression/covariates, measurement error/within‐species variation, compatibility with tip‐dated/non‐ultrametric trees, and software language). Coding reflects documented, user‐facing software functionality (✓ = supported; blank = not supported/typical; ∼ = limited, indirect, or implementation‐dependent). [1, 18, 20, 24-27, 33, 40, 52, 57-75].

In this paper, we introduce *bifrost* (**B**ranch-level **I**nference **F**ramework for **R**ecognizing **O**ptimal **S**hifts in **T**raits), an R package that implements an end-to-end workflow for estimating cladogenic shifts in the evolutionary dynamics of multivariate traits across phylogenies. *bifrost* is designed as a regime-shift search framework built around mvMORPH::mvgls [1, 28]. mvgls provides a penalized-likelihood (PL, *n* < *p*) or likelihood (LL, *n* > *p*) multivariate GLS estimation engine, and *bifrost* provides methods for: 1) candidate regime generation, 2) parallel model fitting, 3) greedy information-criterion-guided shift discovery, 4) per-shift support diagnostics, and 5) user-facing utilities for extracting and visualizing the results. *bifrost v0*.*1*.*3* infers shifts under a multi-rate multivariate Brownian motion model in which regimes differ by proportional scaling of a shared evolutionary variance-covariance matrix (i.e., proportional matrices in the “Flury hierarchy” [29, 30]). This parameterization (the ‘BMM’ model in mvgls) reduces regime complexity to one scalar parameter per regime while simultaneously accounting for multivariate covariance structure (see below).

When other available software approaches are compared across core design features (Table 1), including the target of heterogeneity (e.g., rate, optimum, covariance), inference approach, multivariate scope, and support for regression and error estimation, *bifrost* occupies a distinct region of method-space, bridging high-dimensional multivariate GLS frameworks and automated regime-discovery approaches. Here, we describe the *bifrost* R package, focusing on the model and search algorithm at a level required to understand the software implementation. We then present a worked example using high-dimensional fossil jaw-shape data and discuss how inferred shifts may be translated into testable hypotheses in downstream analyses.

## Materials and Methods

### Model and Parameter Estimation

*bifrost 0*.*1*.*3* automates the discovery of discrete-regime multivariate Brownian motion (mvBM) models that can provide a significantly better fit to a dataset than a uniform Brownian process (see *Shift Search Algorithm* below). Let **Y** be *n* × *p* an matrix of trait values for *n* taxa and *p* traits. Under mvBM, traits evolve along a phylogeny as a multivariate normal process with covariance proportional to shared ancestry, such that the expected covariance among taxa is given by the Kronecker product of the phylogenetic variance–covariance matrix (derived from branch lengths) and an evolutionary variance–covariance matrix Σ describing rates and covariation among traits [1, 4]. In a multi-regime model, branches are assigned to discrete regimes *r* = 0,…,*R*, and each regime is associated with an evolutionary variance–covariance matrix Σ_*r*_ [20, 28, 31, 32].

To enable scalable, multivariate, and time-heterogeneous inference, *bifrost* relies on the scalar rate parameterization implemented in the scalar “BMM” model in mvMORPH::mvgls (multivariate generalized least squares). Under this model, regimes share a common covariance structure but differ in scale, such that Σ_*r*_ = *ρ*_*r*_Σ_0_, where Σ_0_ is a shared baseline covariance “shape” matrix and *ρ*_*r*_ > 0 is a regime-specific scalar. Under both likelihood and penalized-likelihood estimation, the regime parameters returned by mvgls are interpreted as average evolutionary rates across traits within each regime [1]. More generally, mvgls supports a formula-based GLS mean structure, allowing *bifrost* to be used in broad mvPGLS-like settings where hidden rate regimes can influence residual covariance, conceptually similar to other hidden-rates frameworks [33, 34]. In this regression setting, observed predictors enter the GLS mean structure, whereas inferred regimes describe hidden branch-specific residual rate heterogeneity rather than state-dependent rate effects [35]. To our knowledge, *bifrost* provides a unique integrated workflow capable of automatically inferring hidden evolutionary rate regimes from high-dimensional or multivariate datasets.

### Shift Search Algorithm

In a typical analysis, our procedure first proposes candidate shift points at internal nodes of a user-provided phylogeny. Only nodes with descendant clades that contain at least a user-defined minimum size threshold (min_descendant_tips) of extant or fossil taxa are considered, which (1) reduces the model search space (which otherwise scales as 2^n^) and (2) helps prevent overfitting shifts to very small clades [21, 36, 37]. Candidate evaluation is thus primarily a function of tree size and this user-defined constraint, which must be ≥ 2.

Shift discovery proceeds in two stages (Figure 1). First, *bifrost* fits a single-regime global baseline model and then evaluates each candidate node as a one-shift, two-regime model (Figure 1). Candidate two-regime models are compared to the baseline using an information criterion selected by the user: currently, the generalized information criterion (GIC) [38] and the Bayesian information criterion (BIC) [39] are available. All candidate models are then ranked by their improvement in IC relative to the baseline (ΔIC). Second, *bifrost* builds a multi-shift configuration using a greedy, stepwise aggregation procedure (recursive clade partition), adapted from [37, 40] (Figure 1). This algorithm iterates over the ranked candidate nodes, attempting to add each shift to the current best configuration and refitting the aggregated model after each proposed addition. A proposed shift is accepted if the IC improves by at least shift_acceptance_threshold units; otherwise, the shift is rejected, and the search moves to the next candidate node.

**Figure 1.**
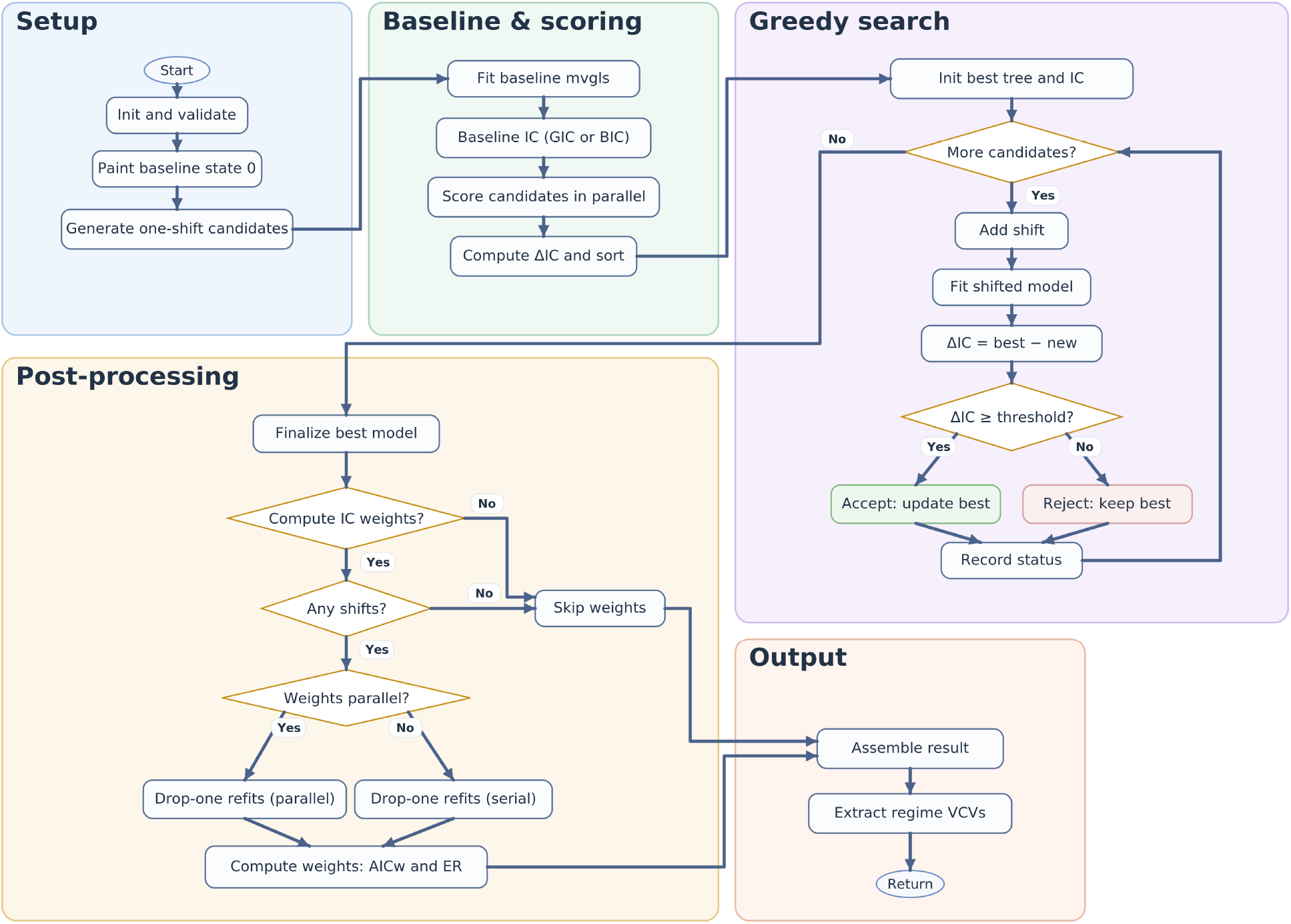
Workflow for multivariate evolutionary regime shift inference in *bifrost*. Starting from a phylogeny and multivariate trait data, *bifrost* first fits a baseline single-regime mvgls model and evaluates all eligible one-shift alternatives using an information criterion (GIC or BIC). Candidate shifts are ranked by ΔIC and then assessed in a greedy stepwise search, in which each proposed shift is added to the current best model and retained only if it improves fit beyond a user-specified threshold. Once no candidates remain, the best-supported multi-regime configuration is finalized. Optional post-processing estimates conditional support for retained shifts via drop-one refits and information-criterion weights. The procedure returns the fitted optimal model, inferred shift nodes, model-comparison summaries, painted regime mappings, and regime-specific variance-covariance matrices.

This strategy yields a practical heuristic search over multi-shift configurations for both large candidate sets (∼10^3^) and high-dimensional datasets (∼10^3-4^). Because the procedure is greedy (e.g., each step attempts to maximize expected improvement in model fit), the resulting configuration is not guaranteed to be globally optimal among all possible shift configurations. Nonetheless, simulations under both generating and misspecified models indicate good performance (see below) [41].

### Per-shift support (IC-weights)

After identifying a final optimized configuration of shifts, *bifrost* can quantify the conditional contribution of each shift using drop-one refits (typically in parallel). For each accepted shift, the corresponding regime is removed from the aggregated configuration, and the reduced model is refit. The information criterion (IC) difference between the full and reduced models is converted to an IC weight *w*, yielding an interpretable per-shift support summary metric conditional on the final model structure [36, 37, 41]:

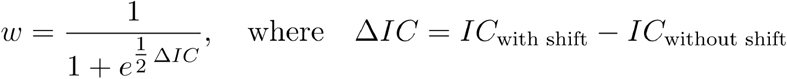

Weights near 1 indicate that removing the shift substantially degrades overall model fit, whereas weights near 0.5 indicate weak or equivocal support. Weights < 0.5 suggest removing the shift improves overall model fit. In practice, however, few shifts are typically detected as low-support when the specified shift_acceptance_threshold is sufficiently large (as shown below).

### Search tuning

Two user-controlled parameters most directly tune the shift search: the minimum descendant-tip filter (min_descendant_tips), which constrains candidate locations, and the ΔIC acceptance threshold (shift_acceptance_threshold), which governs shift acceptance. Larger minimum clade sizes and higher ΔIC thresholds reduce the likelihood of inferring shifts in small clades (where power is low) or shifts that yield only marginal improvements in fit. Because information-criterion behavior depends on dataset size and trait dimensionality, we recommend conservative defaults guided by prior simulations [41], while noting that performance may vary across datasets. In practice, conservative settings, such as ΔGIC = 20 and a minimum clade size of 10, tend to prioritize shifts that yield the largest improvements in model fit and provide good false-positive and false-negative rates with large datasets [41]. Sensitivity analyses can be performed by repeating runs across a range of search parameters, or, alternatively, parametric simulation can be used to identify optimized search parameters suited to the statistical properties of a given dataset [41].

### Package Implementation

*bifrost* is implemented as an open-source R package (≥ GPL-2) [42] that wraps multivariate GLS model fitting [1] in a regime-shift discovery workflow. The package is distributed through CRAN (https://doi.org/10.32614/CRAN.package.bifrost), which provides an archive of released versions (including source and platform-specific binaries), with ongoing development and issue tracking hosted on GitHub. We provide comprehensive documentation, including a pkgdown [43] website (https://jakeberv.com/bifrost) and vignettes (https://jakeberv.com/bifrost/articles/). The source repository (https://github.com/jakeberv/bifrost) includes an extensive suite of unit tests implemented with testthat [44], and continuous integration (CI) workflows that run R CMD check across major platforms (macOS, Linux, and Windows) via GitHub Actions [45], including additional checks using R-hub [46]. Testing coverage is tracked with Codecov (https://codecov.io) [47] and currently reports 100% line coverage on the main branch. This testing and CI setup follows best practices for R package development [48, 49], including automated cross-platform checks and coverage tracking. Key dependencies include ape [50], future [51], mvMORPH [28], phytools [52, 53], viridis [54], and txtplot [55]. R code development was assisted by the GPT-4/4o/5 large language models (OpenAI).

### Input Data

The primary function, searchOptimalConfiguration(), implements the workflow described above, and the package includes helper functions for visualizing search behavior and extracting summary statistics. searchOptimalConfiguration() requires (i) a phylogenetic tree with branch lengths in time units and (ii) a multivariate trait matrix with row names matching the tree’s tip labels. The input tree may include or be composed of fossil tips (mvMORPH::mvgls does not require ultrametricity). Trait data may be any continuous multivariate representation provided as an *n* × *p* matrix aligned to the tree, including Generalized Procrustes Analysis (GPA) landmark coordinates that have been vectorized into two dimensions. Notably, this workflow is not restricted to phenotypic data and is applicable to other continuous multivariate datasets aligned to phylogenies, including suites of ecological or functional traits, environmental niche variables, multivariate life-history data, multivariate genomic characteristics (e.g., gene expression profiles, or regulatory or compositional summaries), or other learned embeddings, provided they can reasonably be treated as continuous traits evolving along a phylogeny under a mixture of Brownian-like covariance models.

Researchers specify a formula string to describe the fixed-effects regression structure. For example, trait_data ∼ 1 is appropriate for GPA coordinate matrices (example below), whereas for log-linear measurements, users could include log-mass as a covariate (e.g., trait_data ∼ log(mass)) to account for size-related allometry under a Brownian covariance structure [41, 56]. Additional or alternative predictor effects are thus estimated jointly under the inferred phylogenetic covariance structure. Users may also pass various model-fitting options through *bifrost* to mvMORPH::mvgls (e.g., estimation method, penalty/target settings, and measurement-error modeling).

### Outputs

searchOptimalConfiguration() returns a structured S3 results object (class bifrost_search) containing the recorded user inputs, inferred shift node IDs, baseline and optimal information-criterion values (and ΔIC), the final fitted mvgls object, and a SIMMAP-class [52] tree object containing the optimized regime configuration. If per-shift support is requested, the output includes information-criterion weights for each accepted shift. When store_model_fit_history = TRUE, *bifrost* saves the full model search history, including intermediate mvgls model fit objects and whether candidate shifts were accepted or rejected at each step. To limit memory overhead during the search, intermediate history files are written to a temporary directory and subsequently assembled into the results object. The output object is designed for downstream plotting, summaries, and various post-hoc analyses, including the visualization of regime mappings and the inspection of search history (see discussion below). We also provide a custom print() method that summarizes candidate and shift counts, key tuning settings, core mvgls fit details (model, method, penalty/target, formula), and optional diagnostics such as per-shift weights and a text-based IC history plot.

### Parallelization

The computation that occurs within *bifrost* happens across three stages (Figure 1): (1) candidate generation and scoring (2) sequential greedy aggregation of shifts, and (3) optional post hoc support refits. The first and third stages can be easily parallelized because they involve fitting many independent mvgls models. In the candidate-scoring stage (1), *bifrost* fits a baseline model and a set of one-shift candidate models, each corresponding to a different shift candidate. In the support-estimation stage (3), bifrost refits the final model repeatedly after removing one shift at a time (drop-one refits). In contrast, the greedy aggregation stage (2) is inherently sequential. Each iteration proposes adding a shift to the current best configuration, refits the updated multi-shift model, and then accepts or rejects the proposal based on the ΔIC threshold. Because each step depends on the set of already accepted shifts, this stage cannot be easily parallelized and often dominates the runtime.

Parallel execution is implemented using the future framework [51], with the number of workers set by num_cores. *bifrost* automatically switches to an appropriate future plan within the parallel candidate-scoring and optional drop-one refit stages, and restores a sequential plan afterward. On Unix-like systems where forking is available, *bifrost* uses multicore to reduce overhead; otherwise, it uses multisession (e.g., on Windows PCs). Before launching parallel tasks, *bifrost* caps BLAS/OpenMP-related thread settings to 1 per worker via environment variables to prevent CPU oversubscription, and restores the prior thread settings at the end of the parallel block. For large datasets, researchers may need to increase the future.globals.maxSize option so that required objects can be passed to worker processes.

## Paleozoic fish jaw shape macroevolution

We provide an example use case of *bifrost* using a fossil dataset from [76] that quantifies jaw shape evolution during the early radiation of bony fishes. [76] shows that these early lineages exhibit heterogeneous rates of morphological evolution, with “macroevolutionary role reversals” between sarcopterygian (lobe-finned) and actinopterygian (ray-finned) clades. The example dataset includes a time-scaled phylogeny (86 taxa) and an associated set of Generalized Procrustes–aligned 3D landmark coordinates. Landmark data comprise 126 landmarks per taxon, stored as a 126 × 3 × 86 array of X–Y–Z coordinates (32,508 scalar coordinate values). Example files are bundled with the package under extdata and are also available from the data repository associated with the original study.

### Workflow

This example (adapted from the full vignette on the package website) first loads the phylogeny and landmark data, converts the landmark array into a matrix with rows aligned to the tree-tip order, and then calls searchOptimalConfiguration(). We choose conservative search settings (minimum clade size of 10 and a ΔGIC threshold of 20) to prioritize strongly supported shifts. We also use helper functions from the geomorph and ape [50] R packages [77] to preprocess the dataset into a 2D array (86 × 378). Within *bifrost*, we set the method parameter to use the “H&L” penalized-likelihood method [78]. This method is passed internally to mvMORPH::mvgls to provide a fast, analytical approximation of Leave-One-Out Cross-Validation (LOOCV) for high-dimensional covariance matrix estimation. Other methods are available and described in the mvMORPH package documentation [1, 28].

Runtime for datasets at the scale of the provided example (e.g., 86 terminals, below) is modest (e.g., < 15 minutes on an M1 Max MacBook Pro). However, these times can be significantly extended, depending on the parameter estimation strategy and tree size. While the example we provide here is relatively lightweight, larger-scale analyses (e.g., [41]) may require significantly more computing time. Runtimes are primarily a function of tree size and the parameter-estimation approach; the best estimation strategy will depend on the analytical scale and the statistical properties of individual datasets.

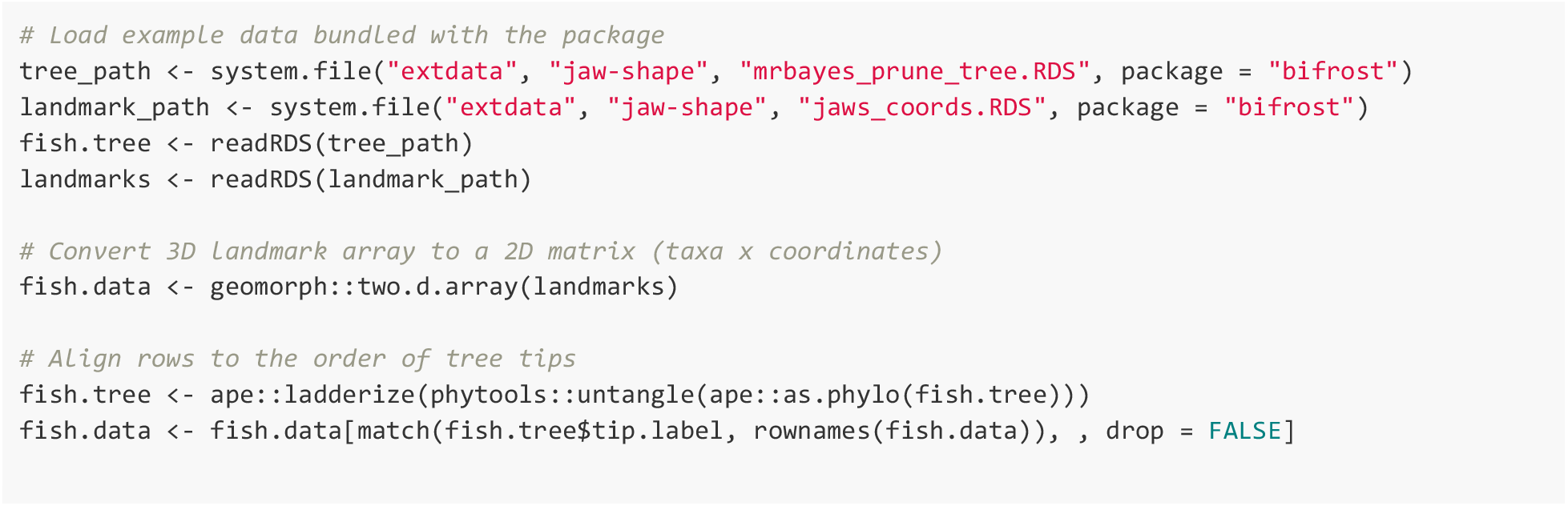
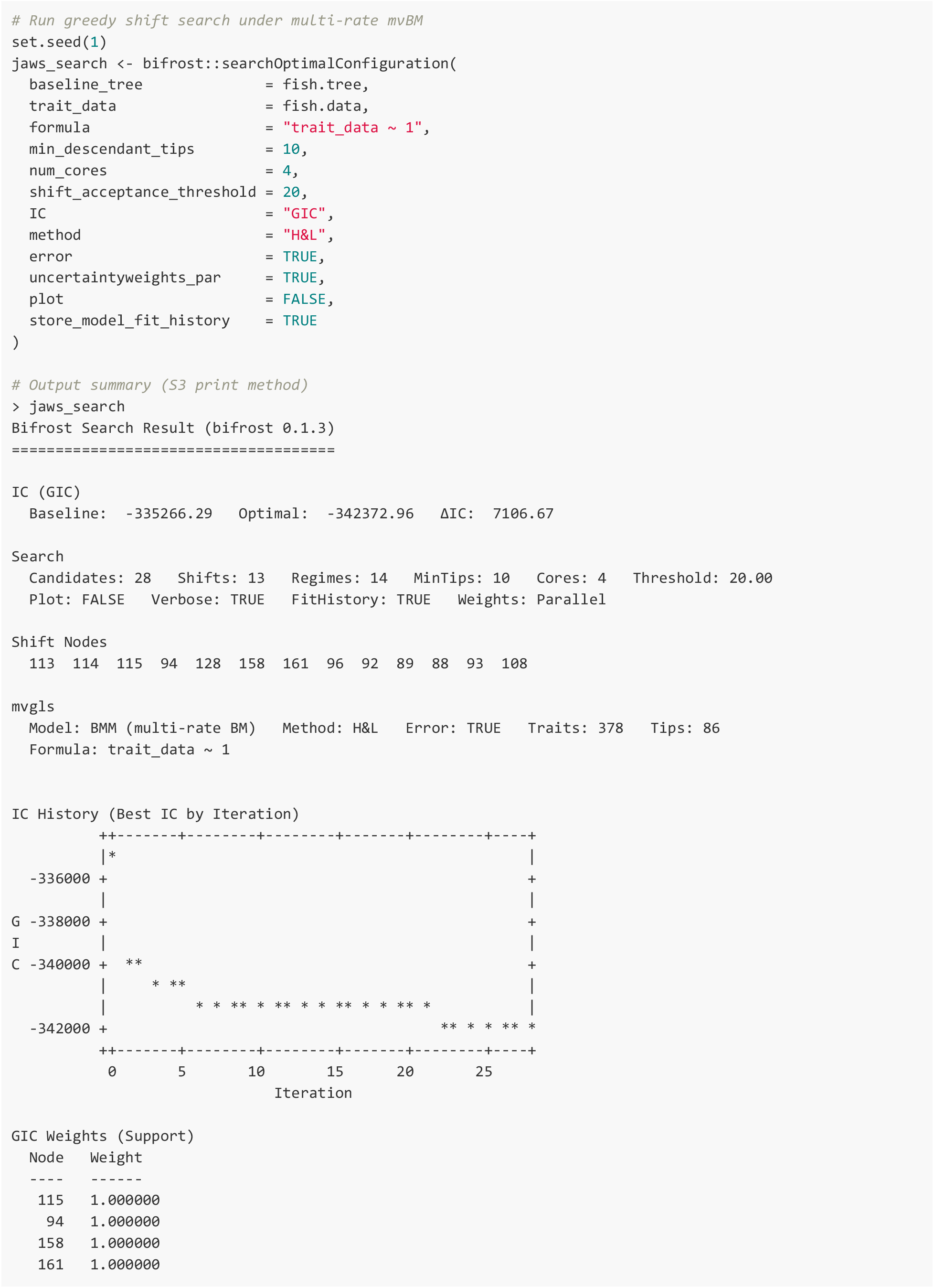
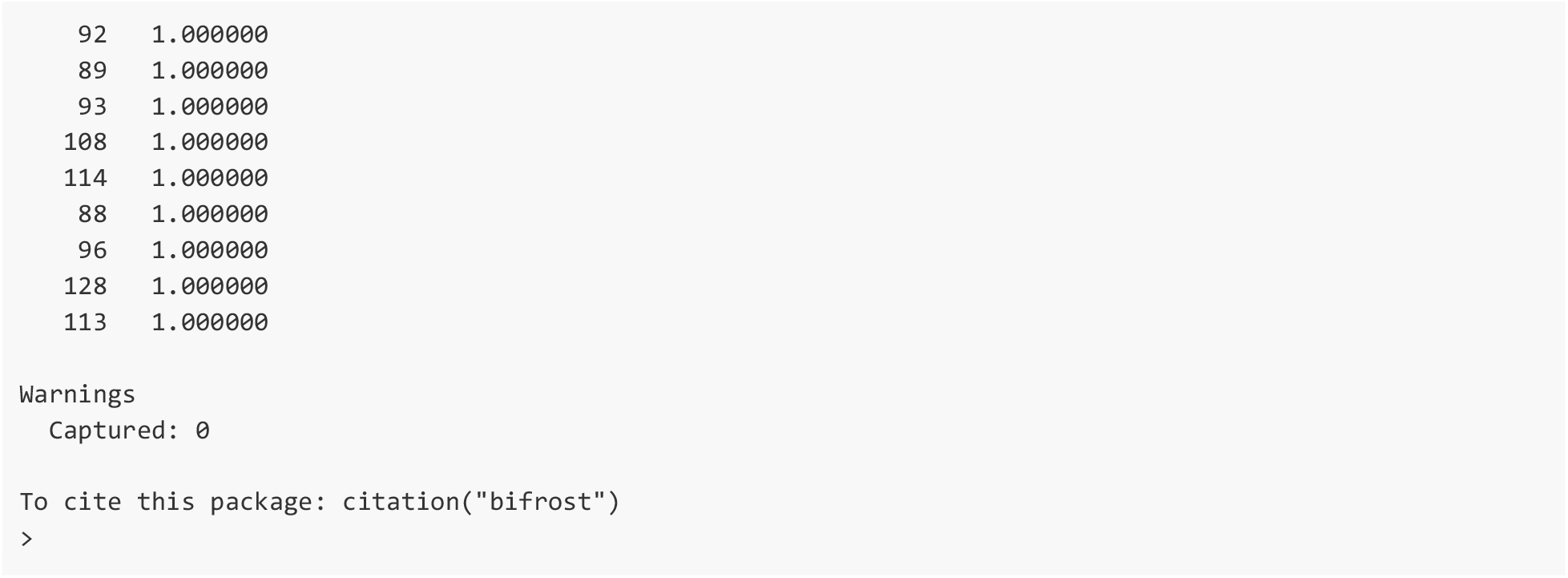

### Interpreting outputs

*bifrost* returns an object with class bifrost_search (a structured list object) that includes node identifiers for accepted shifts, the information criterion values for the baseline and optimal models, the final ‘painted’ SIMMAP tree [52], and regime-specific evolutionary variance-covariance estimates. Under the multi-rate Brownian Motion (BMM) model used here, regimes share a common covariance structure; thus, the returned VCV matrices serve primarily as diagnostic outputs. When per-shift support is requested, IC weights provide a simple diagnostic of which inferred shifts are strongly supported conditional on the final configuration.

In the worked example of Paleozoic fish jaw evolution, the inferred multi-regime configuration yielded a GIC improvement of more than 7,000 units relative to the single-rate baseline model. This magnitude of model-fit improvement provides decisive statistical evidence for multivariate rate heterogeneity, supporting the hypothesis that early sarcopterygian and actinopterygian lineages explored morphospace under fundamentally different evolutionary dynamics, in line with the conclusions of [76]. The mapped output phylogeny encodes this heterogeneity as a computable object that identifies lineages in which shifts in evolutionary rates may have occurred (Figure 2).

**Figure 2.**
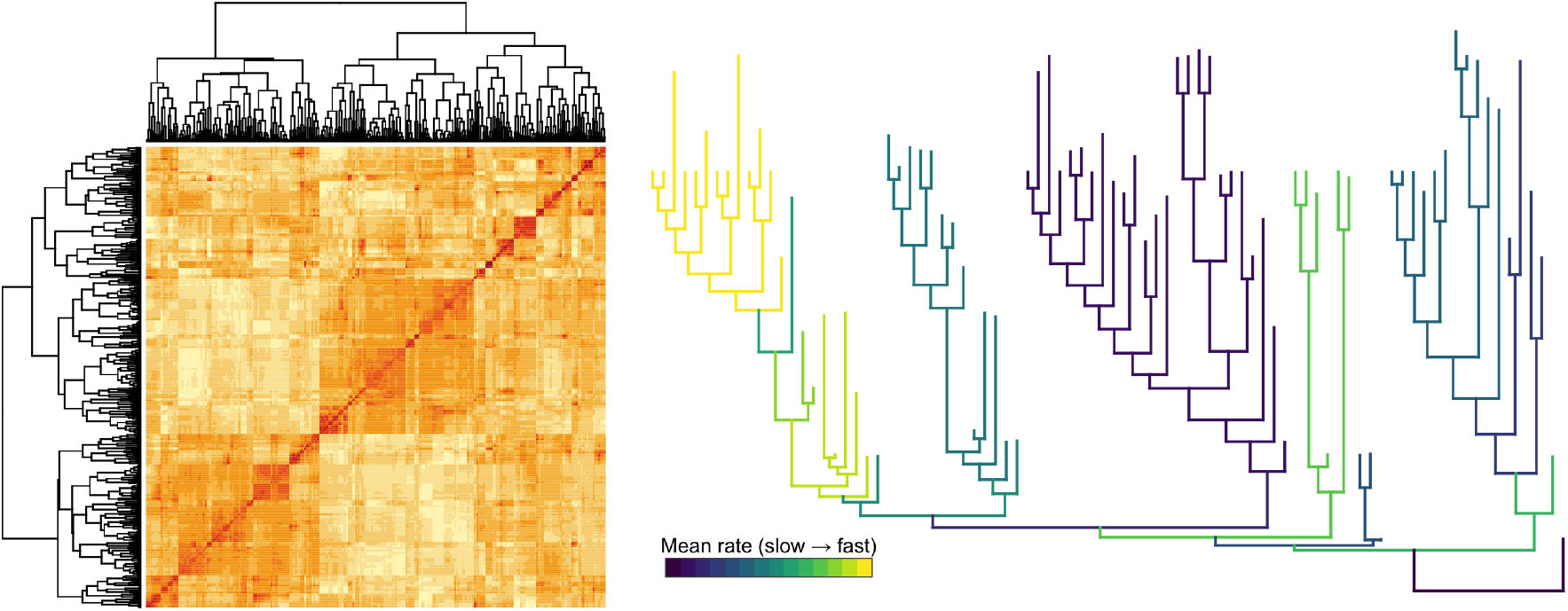
Examples of *bifrost* output visualizations. Left: Heatmap of estimated evolutionary correlations (computed as cov2cor(jaws_search$VCVs$`0`)) among jaw-shape landmarks in the example analysis above (378×378). Rows and columns are ordered by hierarchical clustering, and blocks of high correlation along the diagonal suggest modular patterns of trait covariation. Right: SIMMAP-class phylogeny object with branches colored by estimated mean evolutionary rate under the optimized multivariate multi-regime model. To show overall variation in inferred evolutionary tempo across the phylogeny of Paleozoic fishes, branches are colored using a viridis palette [54], with yellow indicating faster rates and dark purple indicating slower rates. Faster branches on the left side correspond to lungfishes; see [76] for further discussion.

## Discussion

*bifrost* addresses a growing need for phylogenetic comparative methods that operate on multivariate or high-dimensional trait datasets. It provides an explicit, reproducible framework for discovering rate regimes under a multivariate Brownian motion model while operating directly in trait space. By combining multivariate GLS estimation with an automated regime-shift search, *bifrost* enables researchers to move from raw multivariate trait data and a phylogeny to an interpretable set of supported shifts, regime-specific evolutionary covariance estimates, and diagnostic summaries. Our strategy is deliberately pragmatic: it uses a deterministic model-selection algorithm, parallel candidate evaluation, and user-controlled acceptance thresholds to produce a single well-supported configuration that is straightforward to interpret and visualize.

A key motivation for operating directly in trait space is to avoid distortions in evolutionary inference that can arise from dimensionality reduction. As highlighted by Uyeda et al. [14], representing multivariate phenotypes with a small subset of principal components can discard multivariate structure relevant to evolutionary processes. Furthermore, when the retained components are modeled independently, early components may artifactually resemble an “early burst” dynamic [79, 80], whereas later components can resemble stabilizing selection typical of an OU process. These distortions arise because PCA maximizes total trait variance without accounting for phylogenetic relationships, effectively sorting axes by statistical variance rather than phylogenetically relevant signal [13, 14]. As a result, typical PCA-based workflows may filter out dimensions that account for little overall variance but nonetheless may contribute meaningfully to multivariate evolutionary patterns [10, 14]. Phylogenetically informed PCA (e.g., [56]) can mitigate some of these issues [14], but ordination still imposes a model-dependent projection that can obscure multivariate structure.

In this context, *bifrost**’**s* design choices prioritize scalability and interpretability for regime discovery across high-dimensional datasets. Biologically, *bifrost* focuses on partitioning phylogenetic branches into regimes that differ in multivariate evolutionary rates under a shared covariance structure. The “proportional rate matrix” assumption of *bifrost* dramatically reduces the parameter space compared to fully Bayesian multivariate approaches (e.g., ratematrix [57]). *bifrost* (and by extension, the “BMM” model in mvgls) thus sacrifices some model generality in exchange for scalability—a tradeoff that enables exploration of extremely large datasets. Further, *bifrost* supports phylogenetic regression and measurement error estimation natively through its interface to mvMORPH [1, 28, 81, 82]. This enables researchers to include covariates that naturally account for mean trends in the data (e.g., body mass, trophic niche), which mvgls will jointly estimate alongside rate regimes. In that mode, *bifrost* seeks shifts in a residual evolutionary process, decoupling mean changes from changes in evolutionary volatility or constraint (under multivariate Brownian motion).

*bifrost* has been applied in an empirical analysis of functional traits across >2,000 species [7, 41], and that study included extensive validation of the method, including simulation-based assessments of power, accuracy, and robustness across a broad range of scenarios (e.g., varying numbers and magnitudes of shifts, different trait dimensions, and model misspecification [41]). Under empirically derived simulation conditions, *bifrost* showed high accuracy in identifying true shift locations and essentially zero false-positive rates, with balanced accuracy exceeding 90% [41]. These evaluations indicate that *bifrost* is a statistically robust tool for detecting regime shifts and demonstrate a pipeline for conducting similar validation exercises with other datasets. We encourage researchers to follow these guidelines when evaluating expected model performance for individual datasets, which may differ substantially from the statistical properties examined in that study [41]. In practice, conservative settings (e.g., shift_acceptance_threshold = 20, min_descendant_tips = 10) tend to yield a manageable number of well-supported shifts while guarding against overfitting.

Beyond identifying potential macroevolutionary regime shifts, the output of *bifrost* is structured to facilitate deeper biological analysis, and the *bifrost* package will evolve to include additional tools for downstream analysis. The principal output—a SIMMAP-style phylogeny object [52], annotated with regimes, parameter estimates for each regime, and per-branch regime assignments—enables a wide array of downstream analyses. By flagging specific tree partitions with unusual evolutionary dynamics, *bifrost* supports a “phylogenetic natural history” approach [83] in which inferred shifts serve as testable hypotheses for targeted follow-up analyses with independent ecological, behavioral, environmental, or genomic data.

For example, researchers can examine where and when shifts are detected to investigate whether phenotypic transitions align with shifts toward canalized phenotypic architectures [22] or with geological transitions [36]. More generally, one could use *bifrost* to investigate whether shifts in phenotypic evolution correlate with shifts in behavior (e.g., origin of sociality, migration), shifts in ecological factors (e.g., habitat use or trophic niche), or even genomic traits (such as polyploidy events, changes in developmental gene regulatory networks, or life-history genomic correlates). Digging deeper, one could also use *bifrost* as a starting point for comparing inferred high- and low-rate phenotypic regimes to examine how model shifts relate to patterns of phenotypic integration (e.g., the internal organization of phenotypic covariation). The output of *bifrost* is also inherently a spatio-temporal hypothesis that can be cross-referenced with external geographic data. Looking forward, there are several natural extensions of *bifrost*. Since many biological questions center on putatively directional or adaptive trait changes [19, 31, 84], generalizing our modeling framework beyond Brownian motion to include the multivariate Ornstein–Uhlenbeck (OU) processes [24] would open important avenues of research. Under the scalar OU (scOU) model, the target of variation is the OU optimum (i.e., a shift in a vector of optimum trait values for regime r: *θ*_*r*_). The scOU model assumes a single selection-strength parameter is shared across all traits, while traits may be correlated [24]. Incorporating OU dynamics will allow *bifrost* to detect directional shifts in high-dimensional regimes, as well as patterns of phenotypic convergence (e.g., [26]). The existing PhylogeneticEM [24], ℓ1ou [65], and mvSLOUCH [68] methods represent significant steps in these directions, and future extensions of *bifrost* will expand this body of work by supporting automated inference of directional evolutionary processes directly in high-dimensional trait space. Finally, ongoing work will expand *bifrost**’s* workflow capabilities for integrated post-hoc analyses: for example, additional functions for automated post-hoc analyses of trait covariance structures in inferred regimes, simulations for model performance, and spatial analyses [41].

## Conclusion

In sum, *bifrost* provides a practical and scalable approach for estimating shifts in multivariate trait evolution on phylogenies and offers substantial flexibility through regression-based modeling. By operating directly in trait space, *bifrost* helps preserve the biological structure of high-dimensional datasets when studying evolutionary heterogeneity and supports the development of explicit, testable phylogenetic hypotheses. These interpretable regime mappings provide a flexible starting point for downstream comparative analyses linking shifts in trait evolution to ecological, environmental, and genomic factors. As more researchers embrace high-dimensional trait datasets–from shape coordinates to gene expression profiles–tools like *bifrost* may help unravel the complex tempo and mode of multivariate trait evolution. We envision *bifrost* as a useful tool for comparative biologists to explore the rich tapestry of evolutionary dynamics across the tree of life.

## Acknowledgements

This work was partly supported by the facilities and staff of the University of Michigan Institute for Data Science and AI in Society, the School for Environment and Sustainability, the Department of Ecology and Evolutionary Biology, and the Museum of Paleontology. Initial development of the *bifrost* R package was carried out with support from the Oxford Research Software Engineering Group as part of the Research Software Engineering Workshop, with funding from Schmidt Sciences, LLC, and support from Minjoon Kouh, Jack Leland, Oliver King, and Timothy Poon. We thank Tom Carruthers, Sean McHugh, James Boyko, Shelly Gaynor, Madisyn Guza, and Charlotte Probst for helpful discussions, and extend special thanks to Julien Clavel for early support in understanding the architecture of mvMORPH. The name *bifrost* is inspired by Bifröst, the rainbow bridge of Norse mythology that connects Earth (Midgard) and Asgard within the cosmic structure of Yggdrasil, the Tree of Life, echoing how our framework links observable data to hidden evolutionary shifts across the history of life. BCW and JSB were supported by the David and Lucile Packard Foundation. JSB, NF, MJT, and HLL were each supported by an Eric and Wendy Schmidt AI in Science Postdoctoral Fellowship (Schmidt Sciences, LLC).

## CRediT (Contributor Roles Taxonomy)

Conceptualization: JSB

Methodology: JSB, SAS

Software: JSB, NF, MJT, HLL, SAS

Validation: JSB

Formal analysis: JSB

Investigation: JSB, RARV, EMT

Resources: JSB, SAS, MF, DFF, BCW, EMT, RARV

Data Curation: JSB, EMT, RARV

Writing - Original Draft: JSB

Writing - Review & Editing: JSB, NF, MJT, HLL, EMT, RARV, SAS, MF, DFF, BCW

Visualization: JSB Supervision: JSB, BCW

Project administration: JSB, BCW

Funding acquisition: JSB, NF, MJT, HLL, SAS, DFF, BCW

## Statement on inclusion

Our research brings together authors from multiple institutions and countries to develop open-source software tools that are broadly accessible to the global scientific community. Although the development of this software did not involve field data collection requiring local stakeholder engagement, we have prioritized open science practices by making all code, documentation, and tutorials freely available, thereby lowering barriers to access for researchers worldwide.

## Data Availability

The *bifrost* package is open source and available under a ≥ GPL-2 license. The current release is archived on CRAN (https://doi.org/10.32614/CRAN.package.bifrost), and the development version is hosted on GitHub (https://github.com/jakeberv/bifrost). The empirical fish jaw dataset analyzed in this study is available from the bifrost package (system.file(“extdata”, …)), and the original data are available from [76].

## References

1. Clavel, J., L. Aristide, and H. Morlon, A Penalized Likelihood Framework for High-Dimensional Phylogenetic Comparative Methods and an Application to New-World Monkeys Brain Evolution. Systematic Biology, 2019. 68(1): p. 93–116.

2. He, Y., et al., Opportunities and Challenges in Applying AI to Evolutionary Morphology. Integrative Organismal Biology, 2024. 6(1).

3. Gould, S.J., et al., The spandrels of San Marco and the Panglossian paradigm: a critique of the adaptationist programme. Proceedings of the Royal Society of London. Series B. Biological Sciences, 1979. 205(1161): p. 581–598.

4. Lande, R., Natural selection and random genetic drift in phenotypic evolution. Evolution, 1976. 30(2): p. 314–334.

5. Felsenstein, J., Phylogenies and quantitative characters. Annual Review of Ecology and Systematics, 1988. 19: p. 445–471.

6. Mitteroecker, P. and F. Bookstein, Linear Discrimination, Ordination, and the Visualization of Selection Gradients in Modern Morphometrics. Evolutionary Biology, 2011. 38(1): p. 100–114.

7. Weeks, B.C., et al., Skeletal trait measurements for thousands of bird species. Scientific Data, 2025. 12(1): p. 884.

8. Weeks, B.C., et al., A deep neural network for high-throughput measurement of functional traits on museum skeletal specimens. Methods in Ecology and Evolution, 2023. 14(2): p. 347–359.

9. Weaver, W.N. and S.A. Smith, From leaves to labels: Building modular machine learning networks for rapid herbarium specimen analysis with LeafMachine2. Applications in Plant Sciences, 2023. 11(5): p. e11548.

10. Adams, D.C. and M.L. Collyer, Multivariate Phylogenetic Comparative Methods: Evaluations, Comparisons, and Recommendations. Systematic Biology, 2017. 67(1): p. 14–31.

11. Adams, D.C., A Generalized K Statistic for Estimating Phylogenetic Signal from Shape and Other High-Dimensional Multivariate Data. Systematic Biology, 2014. 63(5): p. 685–697.

12. Collyer, M.L., E.K. Baken, and D.C. Adams, A standardized effect size for evaluating and comparing the strength of phylogenetic signal. Methods in Ecology and Evolution, 2022. 13(2): p. 367–382.

13. Bookstein, F.L., Random walk as a null model for high-dimensional morphometrics of fossil series: geometrical considerations. Paleobiology, 2013. 39(1): p. 52–74.

14. Uyeda, J.C., D.S. Caetano, and M.W. Pennell, Comparative Analysis of Principal Components Can be Misleading. Systematic Biology, 2015. 64(4): p. 677–689.

15. LeCun, Y., Y. Bengio, and G. Hinton, Deep learning. Nature, 2015. 521(7553): p. 436–444.

16. Vaswani, A., et al., Attention is all you need. Advances in neural information processing systems, 2017. 30.

17. Dzieżyc, M., et al., Can we ditch feature engineering? end-to-end deep learning for affect recognition from physiological sensor data. Sensors, 2020. 20(22): p. 6535.

18. Rabosky, D.L., Automatic Detection of Key Innovations, Rate Shifts, and Diversity-Dependence on Phylogenetic Trees. PLOS ONE, 2014. 9(2): p. e89543.

19. Simpson, G.G., Tempo and mode in evolution. 1944: Columbia University Press.

20. O’Meara, B.C., et al., Testing for different rates of continuous trait evolution using likelihood. Evolution, 2006. 60(5): p. 922–933.

21. Parins-Fukuchi, C.T., et al., Transitions Into Freezing Environments Linked With Shifts in Phylogenetic Integration Between Vitaceae Leaf Traits. Ecology and Evolution, 2024. 14(11): p. e70553.

22. Wagner, P.J., Early bursts of disparity and the reorganization of character integration. Proceedings of the Royal Society B: Biological Sciences, 2018. 285(1891): p. 20181604.

23. Pagel, M., C. O’Donovan, and A. Meade, General statistical model shows that macroevolutionary patterns and processes are consistent with Darwinian gradualism. Nature Communications, 2022. 13(1): p. 1113.

24. Bastide, P., et al., Inference of Adaptive Shifts for Multivariate Correlated Traits. Systematic Biology, 2018. 67(4): p. 662–680.

25. Eastman, J.M., et al., A novel comparative method for identifying shifts in the rate of character evolution on trees. Evolution, 2011. 65(12): p. 3578–3589.

26. Ingram, T. and D.L. Mahler, SURFACE: detecting convergent evolution from comparative data by fitting Ornstein-Uhlenbeck models with stepwise Akaike Information Criterion. Methods in Ecology and Evolution, 2013. 4(5): p. 416–425.

27. Uyeda, J.C. and L.J. Harmon, A Novel Bayesian Method for Inferring and Interpreting the Dynamics of Adaptive Landscapes from Phylogenetic Comparative Data. Systematic Biology, 2014. 63(6): p. 902–918.

28. Clavel, J., G. Escarguel, and G. Merceron, mvmorph: an r package for fitting multivariate evolutionary models to morphometric data. Methods in Ecology and Evolution, 2015. 6(11): p. 1311–1319.

29. Phillips, P.C. and S.J. Arnold, Hierarchical comparison of genetic variance-covariance matrices. I. Using the Flury hierarchy. Evolution, 1999. 53(5): p. 1506–1515.

30. Stepanova, N., et al., Punctuated Versus Gradual Shifts in the Multivariate Evolutionary Process: A Test With Paired Radiations of Scincid Lizards. Systematic Biology, 2025. 74(1): p. 1–17.

31. Butler, Marguerite A. and Aaron A. King, Phylogenetic Comparative Analysis: A Modeling Approach for Adaptive Evolution. The American Naturalist, 2004. 164(6): p. 683–695.

32. Revell, L.J. and D.C. Collar, PHYLOGENETIC ANALYSIS OF THE EVOLUTIONARY CORRELATION USING LIKELIHOOD. Evolution, 2009. 63(4): p. 1090–1100.

33. Boyko, J.D., B.C. O’Meara, and J.M. Beaulieu, A novel method for jointly modeling the evolution of discrete and continuous traits. Evolution, 2023. 77(3): p. 836–851.

34. Beaulieu, J.M., B.C. O’Meara, and M.J. Donoghue, Identifying Hidden Rate Changes in the Evolution of a Binary Morphological Character: The Evolution of Plant Habit in Campanulid Angiosperms. Systematic Biology, 2013. 62(5): p. 725–737.

35. May, M.R. and B.R. Moore, A Bayesian Approach for Inferring the Impact of a Discrete Character on Rates of Continuous-Character Evolution in the Presence of Background-Rate Variation. Systematic Biology, 2019. 69(3): p. 530–544.

36. Berv, J.S., et al., Genome and life-history evolution link bird diversification to the end-Cretaceous mass extinction. Science Advances, 2024. 10(31): p. eadp0114.

37. Smith, S.A., N. Walker-Hale, and C.T. Parins-Fukuchi, Compositional shifts associated with major evolutionary transitions in plants. New Phytologist, 2023. 239(6): p. 2404–2415.

38. Konishi, S. and G. Kitagawa, Generalised Information Criteria in Model Selection. Biometrika, 1996. 83(4): p. 875–890.

39. Schwarz, G., Estimating the Dimension of a Model. The Annals of Statistics, 1978. 6(2): p. 461–464, 4.

40. Mitov, V., K. Bartoszek, and T. Stadler, Automatic generation of evolutionary hypotheses using mixed Gaussian phylogenetic models. Proceedings of the National Academy of Sciences, 2019. 116(34): p. 16921–16926.

41. Berv, J.S., et al., Rates of passerine body plan evolution in time and space. Nature Ecology & Evolution (in press), 2026.

42. R Core Team, R: A Language and Environment for Statistical Computing. 2025, R Foundation for Statistical Computing: Vienna, Austria.

43. Wickham, H., et al., pkgdown: Make Static HTML Documentation for a Package. 2025, Posit Software, PBC.

44. Wickham, H., testthat: Get Started with Testing. The R Journal, 2011. 3: p. 5–10.

45. GitHub, I., GitHub Actions. 2025, GitHub, Inc.

46. Csárdi, G. and M. Salmon, rhub: Tools for R Package Developers. 2025.

47. Codecov, Codecov Code Coverage. 2025, Sentry.

48. Wickham, H., et al., Package ‘devtools’. R Package, Version, 2022. 2(4).

49. Wickham, H. and J. Bryan, R Packages (2e). 2026.

50. Paradis, E., J. Claude, and K. Strimmer, APE: Analyses of Phylogenetics and Evolution in R language. Bioinformatics, 2004. 20(2): p. 289–290.

51. Bengtsson, H., A Unifying Framework for Parallel and Distributed Processing in R using Futures. The R Journal, 2021. 13(2): p. 208–227.

52. Revell, L.J., phytools: an R package for phylogenetic comparative biology (and other things). Methods in Ecology and Evolution, 2012. 3(2): p. 217–223.

53. Revell, L.J., phytools 2.0: an updated R ecosystem for phylogenetic comparative methods (and other things). PeerJ, 2024. 12: p. e16505.

54. Garnier, S., et al., viridis(Lite) - Colorblind-Friendly Color Maps for R. 2024.

55. Bornkamp, B. and I. Krylov, txtplot: Text Based Plots. 2025, Comprehensive R Archive Network (CRAN).

56. Revell, L.J., Size-correction and principal components for interspecific comparative studies. Evolution, 2009. 63(12): p. 3258–3268.

57. Caetano, D.S. and L.J. Harmon, ratematrix: An R package for studying evolutionary integration among several traits on phylogenetic trees. Methods in Ecology and Evolution, 2017. 8(12): p. 1920–1927.

58. Caetano, D.S. and L.J. Harmon, Estimating Correlated Rates of Trait Evolution with Uncertainty. Systematic Biology, 2018. 68(3): p. 412–429.

59. Pagel, M., A. Meade, and D. Barker, Bayesian Estimation of Ancestral Character States on Phylogenies. Systematic Biology, 2004. 53(5): p. 673–684.

60. Pagel, M. and A. Meade, Trait macroevolution in the presence of covariates. Nature Communications, 2025. 16(1): p. 4555.

61. Castiglione, S., et al., A new method for testing evolutionary rate variation and shifts in phenotypic evolution. Methods in Ecology and Evolution, 2018. 9(4): p. 974–983.

62. Thomas, G.H. and R.P. Freckleton, MOTMOT: models of trait macroevolution on trees. Methods in Ecology and Evolution, 2012. 3(1): p. 145–151.

63. Grundler, M.C., D.L. Rabosky, and F. Zapata, Fast Likelihood Calculations for Automatic Identification of Macroevolutionary Rate Heterogeneity in Continuous and Discrete Traits. Systematic Biology, 2022. 71(6): p. 1307–1318.

64. Martin, B.S., et al., Modeling the Evolution of Rates of Continuous Trait Evolution. Systematic Biology, 2022. 72(3): p. 590–605.

65. Khabbazian, M., et al., Fast and accurate detection of evolutionary shifts in Ornstein– Uhlenbeck models. Methods in Ecology and Evolution, 2016. 7(7): p. 811–824.

66. Beaulieu, J.M., et al., Modeling stabilizing selection: expanding the Ornstein-Uhlenbeck model of adaptive evolution. Evolution, 2012. 66(8): p. 2369–2383.

67. Hansen, T.F., J. Pienaar, and S.H. Orzack, A COMPARATIVE METHOD FOR STUDYING ADAPTATION TO A RANDOMLY EVOLVING ENVIRONMENT. Evolution, 2008. 62(8): p. 1965–1977.

68. Bartoszek, K., et al., Fast mvSLOUCH: Multivariate Ornstein–Uhlenbeck-based models of trait evolution on large phylogenies. Methods in Ecology and Evolution, 2024. 15(9): p. 1507–1515.

69. Bartoszek, K., et al., A phylogenetic comparative method for studying multivariate adaptation. Journal of Theoretical Biology, 2012. 314: p. 204–215.

70. Grabowski, M., Blouch: Bayesian Linear Ornstein-Uhlenbeck Models for Comparative Hypotheses. Systematic Biology, 2024. 73(6): p. 1038–1050.

71. Adams, D.C. and M.L. Collyer, Extending phylogenetic regression models for comparing within-species patterns across the tree of life. Methods in Ecology and Evolution, 2024. 15(12): p. 2234–2246.

72. Gaboriau, T., et al., A multi-platform package for the analysis of intra- and interspecific trait evolution. Methods in Ecology and Evolution, 2020. 11(11): p. 1439–1447.

73. Kostikova, A., et al., Bridging Inter- and Intraspecific Trait Evolution with a Hierarchical Bayesian Approach. Systematic Biology, 2016. 65(3): p. 417–431.

74. Elliot, M.G. and A.Ø. Mooers, Inferring ancestral states without assuming neutrality or gradualism using a stable model of continuous character evolution. BMC Evolutionary Biology, 2014. 14(1): p. 226.

75. Landis, M.J., J.G. Schraiber, and M. Liang, Phylogenetic Analysis Using Lévy Processes: Finding Jumps in the Evolution of Continuous Traits. Systematic Biology, 2013. 62(2): p. 193–204.

76. Troyer, E.M., et al., Macroevolutionary role reversals in the earliest radiation of bony fishes. Current Biology, 2025. 35(19): p. 4631–4641.e3.

77. Adams, D.C. and E. Otárola-Castillo, geomorph: an r package for the collection and analysis of geometric morphometric shape data. Methods in Ecology and Evolution, 2013. 4(4): p. 393–399.

78. Hoffbeck, J.P. and D.A. Landgrebe, Covariance matrix estimation and classification with limited training data. IEEE Transactions on Pattern Analysis and Machine Intelligence, 1996. 18(7): p. 763–767.

79. Blomberg, S.P., T. Garland, JR., and A.R. Ives, Testing for phylogenetic signal in comparative data: behavioral traits are more labile. Evolution, 2003. 57(4): p. 717–745.

80. Harmon, L.J., et al., Early bursts of body size and shape evolution are rare in comparative data. Evolution, 2010. 64(8): p. 2385–2396.

81. Beaulieu, J.M. and B.C. O’Meara, Navigating “tip fog”: embracing uncertainty in tip measurements. Evolution, 2025. 79(7): p. 1131–1142.

82. Clavel, J. and H. Morlon, Reliable Phylogenetic Regressions for Multivariate Comparative Data: Illustration with the MANOVA and Application to the Effect of Diet on Mandible Morphology in Phyllostomid Bats. Systematic Biology, 2020. 69(5): p. 927–943.

83. Uyeda, J.C., R. Zenil-Ferguson, and M.W. Pennell, Rethinking phylogenetic comparative methods. Systematic Biology, 2018. 67(6): p. 1091–1109.

84. Hansen, T.F., Stabilizing Selection and the Comparative Analysis of Adaptation. Evolution, 1997. 51(5): p. 1341–1351.

